# Single-cell transcriptomic and TCR analysis of human Cytomegalovirus (hCMV)-specific memory T cells reveals effector and pre-effectors of CD8^+^- and CD4^+^-cytotoxic T cells

**DOI:** 10.1101/2023.06.02.543443

**Authors:** Raunak Kar, Somdeb Chattopadhyay, Anjali Sharma, Kirti Sharma, Shreya Sinha, Gopalakrishnan Aneeshkumar Arimbasseri, Veena S. Patil

## Abstract

**Background:** Latent human Cytomegalovirus (hCMV) infection can pose a serious threat of reactivation and disease occurrence in immune-compromised individuals. Though, T cells are at the core of the protective immune response to hCMV infection, a detailed characterization of different T cell subsets involved in hCMV immunity is lacking.

**Results:** Here, in an unbiased manner, we characterized over 8000 hCMV-reactive peripheral memory T cells isolated from seropositive human donors, at a single-cell resolution by analyzing their single-cell transcriptomes paired with the T cell antigen receptor (TCR) repertoires. The hCMV-reactive T cells were highly heterogeneous and consisted of different developmental and functional memory T cell subsets such as, long-term memory precursors and effectors, T helper-17, T regulatory cells (T_REGs_) and cytotoxic T lymphocytes (CTLs) of both CD4 and CD8 origin. The hCMV-specific T_REGs_, in addition to being enriched for molecules known for their suppressive functions, showed enrichment for the interferon response signature gene sets. The hCMV-specific CTLs were of two types, the pre-effector and effector-like. The co-clustering of hCMV-specific CD4-CTLs and CD8-CTLs in both pre-effector as well as effector clusters suggest shared transcriptomic signatures between them. The huge TCR clonal expansion of cytotoxic clusters suggest a dominant role in protective immune response to CMV.

**Conclusions:** The study uncovers the heterogeneity in the hCMV-specific memory T cells reveling many functional subsets with potential implications in better understanding of hCMV-specific T cell immunity. The data presented can serve as a knowledge base for designing vaccines and therapeutics.

## Background

The acquisition of immunological memory to infections is the hallmark of protective immune response. During this conserved process of T cell immunological memory development, the naive T cells that have not previously encountered antigen, differentiate during the primary infection, into memory T cells that have specialized functions in immune defence to a subsequent infection with the same pathogen. A small number of antigen-specific memory T cells can have an enormous impact in directing or misdirecting an immune response upon secondary infection. However, the rarity of antigen-specific memory T cells have limited the detailed characterization of these cells until very recently. The utilization of single-cell genomics to dissect the heterogeneity and understand th global gene expression patterns of these rare antigen-specific memory T cells has advanced our understanding of various disease-specific T cell memory repertoire in an unbiased manner [1–7].

CD4^+^ memory T cells are classified based on their functional properties and in addition more recently also based on their transcriptomes, into multiple subsets such as but not limited to, T helper (T_H_)1, T_H_2, T_H_17, T_H_1/17, T follicular helper (T_FH_), and T regulatory (T_REG_) cells [8–12]. For example, T_FH_ cells are the specialized providers of help to B cells, and T_REGs_, the regulators of the immune response with their immunosuppressive properties [11, 13, 14]. Furthermore, each of those subtypes can exhibit great functional diversity and plasticity [12]. The peripheral CD4^+^ as well as CD8^+^ memory T cells can also be classified based on their developmental stages as long-lived precursor memory and short-lived effector memory T cells such as, stem cell memory T cells (T_SCM_), central memory T cells (T_CM_), and effector memory T cells (T_EM_), effector memory T cells expressing CD45RA (T_EMRA_) [15]. The CD8^+^ T cells are known for their cytotoxic functions, and the CD4^+^ T cells, predominantly, for their helper function where they provide help to other immune cell types including the CD8^+^ T cells in resolving the infection. However, others and we have shown that CD4^+^ T cells can also function as cytotoxic T cells and are enriched in the CD4-T_EMRA_ compartment and show marked T cell antigen receptor (TCR) clonal expansion similar to the CD8-T_EMRA_ cells in many acute and chronic viral infections including the cytomegalovirus (CMV) [1, 3, 4, 6, 16–22].

Following initial infection, the human CMV (hCMV) establishes persistence or latency in immune-competent individuals, although intermittent viral shedding can occur [23]. In the absence of an effective immune response especially in immune-compromised individuals, the virus can reactivate and cause the disease. Hence, reactivation of hCMV poses a higher risk for individuals undergoing transplant or chemotherapy, immune-compromised Human Immunodeficiency Virus (HIV) infected individuals, and new-borns of hCMV seropositive mothers [23]. Both CD4^+^ and CD8^+^ T cells have been shown to play important roles in the immune response to CMV [24, 25]. While hCMV-specific CD8^+^ T cells have extensively been studied, studies on CD4^+^ T cells are sparse [2]. Considering that the emergence of the hCMV-specific CD4^+^ T cells precedes that of CD8^+^ T cells, they were mostly thought to perform supportive roles by enhancing the CD8^+^ T cell responses to the hCMV infection. However, subsequent studies have demonstrated additional roles of CD4+ T cells, other than the supportive one towards CD8^+^ T cell, in CMV immunity (reviewed in [2]). The poorer or lack of hCMV-specific CD4^+^ T cell responses have been associated with persistent viral shedding in urine and saliva as well as severe disease outcomes [24–27]. In individuals undergoing stem cell or organ transplantation, hCMV-specific CD4^+^ and CD8^+^ T cells have been shown to abrogate reactivation [25–29]. Further, during the primary infection, the role of CD4^+^ T cells has been established in resolution of the disease symptoms [30, 31]. The identification of hCMV-specific T cells has classically relied on the measurement of intracellular cytokine production, predominantly interferon gamma (IFNγ) and to a lesser extent interleukin 10 (IL10), in response to stimulation with peptides derived from various hCMV ORFs (Open Reading Frame) or total viral lysate [2, 6, 16, 32–35]. Several studies, in healthy seropositive individuals, have identified a higher percentage of hCMV-specific T cells, reaching almost as high as ∼5-10% of total CD4^+^ and CD8^+^ T cells for epitopes derived from pp65 and IE-1 proteins of hCMV, with majority of them showing terminally differentiated effector memory phenotype [16, 33, 34]. In addition to producing IFNγ, the hCMV-reactive CD4^+^ T cells have been shown to make cytolytic molecules such as Granzymes, Perforin, and CD107a (degranulation marker), and showed efficient lysis of hCMV infected dendritic cells and controlled virus dissemination *in vitro*, suggesting prominent role of cytotoxic CD4^+^ T cells in hCMV immunity [16–20]. CD4^+^ T cells expressing CD25 and FOXP3 (T_REGs_), producing IL10, an immunosuppressive cytokine, have also been observed and are implicated in immune suppression in CMV infection and reactivation [36–39]. However, a detailed characterization of the spectrum of hCMV-specific CD4^+^ memory T cells with respect to their heterogeneity and molecular signatures have been lacking. Hence, the identification and characterization of the overall hCMV-specific memory T cells with respect to their molecular signatures, in CMV seropositive individuals who are otherwise healthy, will enable the identification of correlates of protective immune response to hCMV.

Hence, in this study, to isolate the spectrum of hCMV-specific T cells, independent of their cytokine profile and Human Leukocyte Antigen (HLA) restriction or not limited to a single peptide (peptide-MHC multimers), we used ARTE (Antigen-Reactive T cell Enrichment) assay [40, 41]. The ARTE assay utilizes the surface expression of co-stimulatory molecules (CD154 and CD137) upon TCR engagement with cognate peptide-MHC (Major Histocompatibility Complex) complex [40, 41]. To dissect the heterogeneity across T cell memory subsets and TCR clonal diversity of hCMV-specific memory T cells, we isolated and performed single-cell transcriptomics analysis of over 8000 hCMV-specific memory T cells, isolated based on the surface expression of CD154^+^ and/or CD137^+^ across 10 seropositive donors, after stimulation with pool of hCMV peptides derived from T cell immuno-dominant epitopes of pp65 and IE-1 proteins. Unbiased clustering based on single-cell transcriptomes identified nine distinct clusters of various functional subsets including, T_REGs_, T_H_17, T_H_1 and T_H_1/17, and cytotoxic CD4^+^ T cells (CD4-CTLs). The hCMV-antigen specific T_REGs_ were enriched for molecules linked to their suppressive function and interferon response genes (T_REG-IFN_). The transcriptomics analysis combined with TCR repertoire analysis revealed that the cytotoxic cells are predominantly of 2 types; pre-effectors and effectors and are highly clonally expanded. Interestingly, we observed that the transcriptomes of pre-effectors and effectors of CD4-CTLs and CD8-CTLs were largely indistinguishable. Together, in an unbiased approach, our study has identified many previously known as well as novel subsets of hCMV-specific memory T cells and their unique transcriptomic signatures, which potentially have implications in understanding the reactivation of hCMV in immune-compromised individuals. This knowledge base can be an important resource for designing vaccines and therapeutics.

## Results

### Single-cell transcriptomic analysis of CMV-specific memory T cells

To identify and understand the heterogeneity and clonal diversity of the spectrum of hCMV-specific T cells across different memory subsets in an unbiased way, we performed single-cell transcriptomic (scRNA-Seq) and single-cell TCR (T Cell antigen Receptor) repertoire analysis (scTCR-Seq) of >8000 hCMV-specific memory T cells (Fig. 1A and B). To isolate the spectrum of hCMV-specific T cells independent of their cytokine profile and HLA restriction, we utilized the widely used ARTE (Antigen-Reactive T cell Enrichment) assay where the surface expression of co-stimulatory markers such as CD154 and CD137 is analysed upon the engagement of T cell antigen receptor (TCR) with cognate pMHC (peptide Major Histocompatibility Complex) complex [1, 3, 7, 40–42]. For single-cell omics experiments, we isolated T cells using florescence activated cell sorting (FACS) based on the surface expression of CD154 and CD137 upon stimulation of PBMCs (Peripheral Blood Mononuclear Cells) from 10 hCMV-seropositive and high peptide responder individuals with peptide pools derived from immunodominant pp65 and IE-1 (Fig. 1A and Additional file 1: Fig. S1A-C). To compare the hCMV-specific memory T cells across known memory subsets, the peptide responding T cells, from different CD4^+^ memory T cell compartments (naïve T cells (T_N_; CD45RA^+^CCR7^+^), central memory T cells (T_CM_; CD45RA^-^CCR7^+^), effector memory T cells (T_EM_; CD45RA^-^ CCR7^-^), and effector memory T cells expressing CD45RA (T_EMRA_; CD45RA^+^CCR7^-^) [15]), were FACS sorted separately and pooled before performing single-cell omics experiments (Fig. 1A and Additional file 1: Fig. S1B and C). Further, considering the enrichment of CD4-CTLs (cytotoxic T lymphocytes) in the CD4-T_EMRA_ subset, to parallelly compare the hCMV-specific CD4-CTLs and CD8-CTLs we also included hCMV-reactive T cells from CD8-CTL compartments (CD8^+^CCR7^-^ (CD8-T_EM_ and -T_EMRA_)) in the single-cell omics experiments (Fig. 1A and Additional file 1: Fig. S1B and C) [4]. An unbiased clustering of 6899 hCMV-specific memory T cells (out of 8062 cells) that passed our quality control parameters (QC; please see methods section), identified 9 distinct clusters based on transcriptomes (Fig. 1B), and the clustering was not influenced by donors or the experimental batch (Additional file 1: Fig. S1D and E; Additional file 2: Table S1). Different developmental T cell memory subsets were distinguished based on the surface expression of oligo-tagged antibodies (antibody derived tags; adt) against CD45RA, CCR7, CD95 and IL7Rα as well as *CCR7*, *CD27*, *CD28* transcripts (Fig. 1C and D; Additional file 2: Table S2) [4, 15]. Transcripts for *CD4* were distributed across all clusters while *CD8B* transcripts were majorly found in clusters 2 and 4 (Fig. 1E). Cells in cluster 2 and 4 were mostly T_EMRA_ cells of both CD4 and CD8 origin as they expressed CD45RA (adt CD45RA) and lacked the expression of CCR7 protein (adt CCR7), as well as transcripts for *CCR7*, *CD28* and *CD27* (Fig. 1C and D) [4, 15]. T_N_ cells were very few and were mostly found in cluster 6 (CD45RA^+^CCR7^+^IL7Rα^+^) (Fig. 1C and D). T_CM_ and T_EM_ cells were distributed across multiple clusters (Clusters 0, 1, 3, 5, 6, 7) (Fig. 1C and D). These observations were independently confirmed by the enrichment of subset-specific gene signatures from T_CM_, T_EM_, and T_EMRA_, as well as common gene signatures from T_CM_-T_EM_, and T_EM_-T_EMRA_ [4] (Fig. 1F; Additional file 1: Fig. S1F; Additional file 2: Table S3). Both CD4- and CD8-T_EMRA_ cells (clusters 2 and 4) majorly showed higher expression of *CD137* while other T_CM_ and T_EM_ clusters (0, 3, 6) showed higher expression of *CD154* (Fig. 1G). Interestingly many of the T cell activation markers showed dynamic expression amongst the clusters; *CD154* and *CD137* showed opposing expression and similar observations were made for *CD69* and *IL2RA* (Fig. 1D and G). Further, co-expression analysis of various T cell activation markers in combination, confirmed the variability in expression (Fig. 1G). Most strikingly, the *CD137*^hi^*CD154*^lo^ cells showed low *CD69* expression, while the *CD154* and *CD137* co-expressing cells expressed similar amounts of *TNFRSF4* transcripts (encoding OX40) (Fig. 1G). Together, these results show that our data well represents various memory subsets that show differential expression of T cell activation markers.

**Fig. 1.**
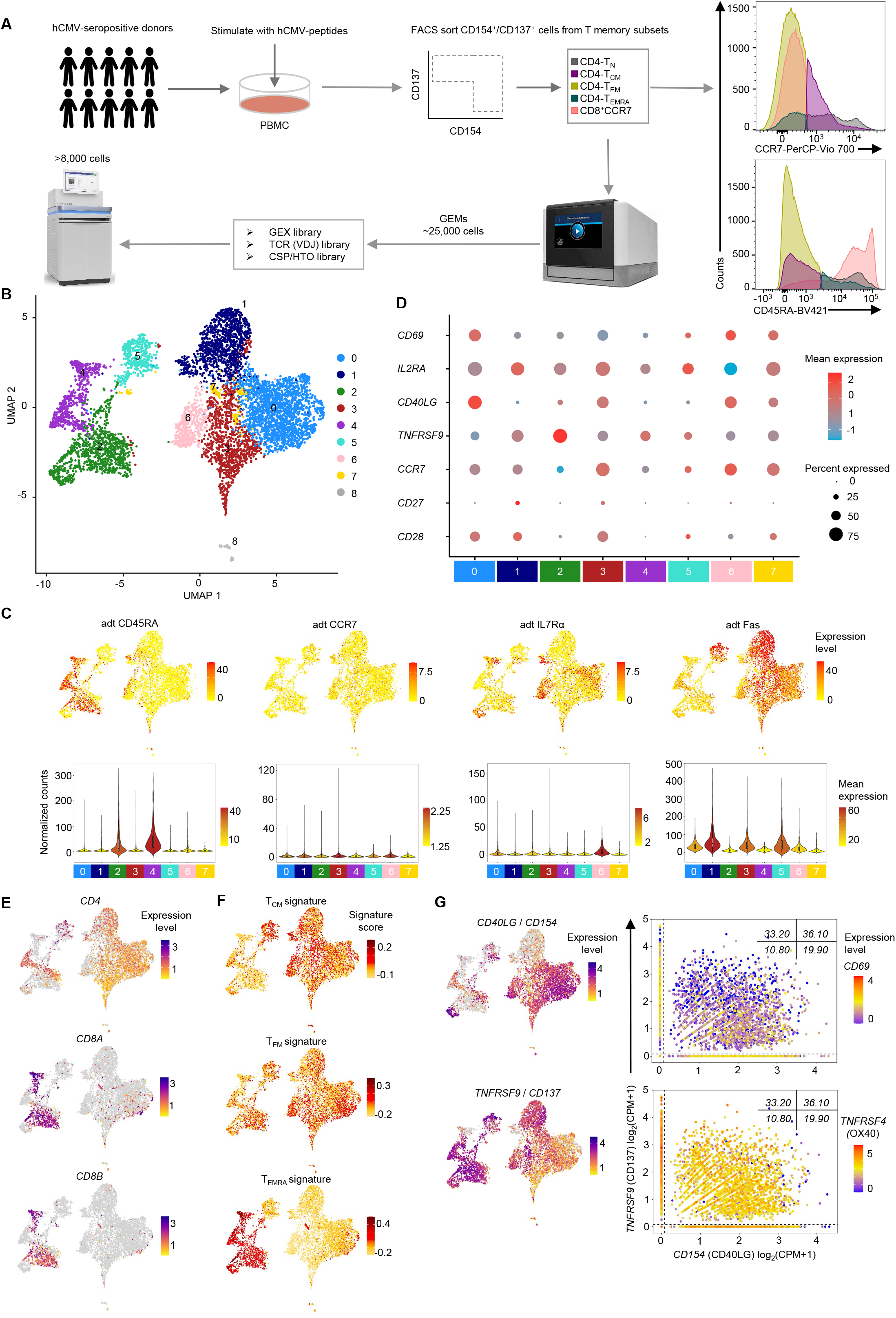
Single-cell transcriptomic analysis of hCMV-reactive memory T cell subsets. (A) Schematic showing the study overview of isolating and single-cell RNA-seq of hCMV-specific T cells using 10X Genomics platform. Histograms (on the right) shows cells isolated using FACS for single-cell RNA-Seq from indicated T cell memory compartment from 10 donors. (B) 2D UMAP of single-cell RNA-Seq data from 6899 cells, using 2000 most variable transcripts, shows 9 distinct clusters of cells of hCMV-specific T cells. (C) 2D UMAP plots (top) and violin plots (bottom) show expression of the indicated protein (oligo-tagged antibodies; adt). Colour scale shows expression levels of individual cells (UMAP) or mean expression in each cluster (violin plot). Only cells (6067) obtained from experiment 2 with 10 donors are shown. (D) Dot plots show the mean expression (colour) and percentage of expressing cells (size) for selected marker gene transcripts in each cluster. (E) 2D UMAP plots of single-cell RNA-Seq data show the expression of the *CD4*, *CD8A* and *CD8B* transcripts. Scale represents expression level of the given transcript. Cells with zero counts for the indicated transcript are shown in grey colour. (F) 2D UMAP plots show the T_CM_, T_EM_ and T_EMRA_ signature score for each cell. (G) 2D UMAP plots (left) of single-cell RNA-Seq data show the expression of the *CD154* (top) and *CD137* (bottom) transcripts. Scale represents expression level of the given transcript. Cells with zero counts for the indicated transcripts are shown in grey colour. Scatter plots (right) show the co-expression matrix for *CD154* (on X-axis), and *CD137* (on Y-axis) with *CD69* or *TNFRSF4* (colour scale: mean expression) transcripts. For violin plots in (C) and dot plots in (D) cluster with less than 1% of the total cells is not shown (cluster 8=56 cells). All expression counts for transcripts are log_2_ normalized (CPM+1).

### hCMV-specific memory T cells are heterogeneous and can be classified into multiple functional T cell subsets

The differential expression analysis of transcripts amongst the nine clusters of hCMV-specific T cells, identified cluster specific expression of transcripts, revealing many functional T cell subsets (Fig. 2A and B; Additional file 2: Table S4). Multiple clusters showed enrichment for long-term memory markers such as *AQP3*, *CCR7*, *TCF7*, *IL7R*, *LTB*, *JUNB, SELL* (Fig. 2A and B) [3, 4]. Cluster 1 showed enrichment for *FOXP3*, *BATF*, *TIGIT* and *TNFRSF1B* transcripts that are known to be expressed in T regulatory cells (T_REG_) (Fig. 2B). Interestingly, cluster 1 also showed enrichment of transcripts encoding interferon (IFN) response signature genes (*ISG20*, *ISG15*, *IFIT3*, *OAS1*, *MX1*) (Fig. 2B). In an independent analysis, cluster 1 cells showed a higher enrichment signature score for overall T_REG_ and IFN response signature gene sets (Fig. 2C; Additional file 2: Table S3), suggesting that cells in cluster 1 are T_REG_ cells and are possibly responding to IFN signalling (T_REG-IFN_). Actively cycling cells were found in cluster 3, one of the clusters enriched for T_CM_ signatures (Fig. 2B and C; Additional file 2: Table S3) [43]. Cells in clusters 2 and 4, enriched for T_EMRA_-specific gene sets, expressed transcripts encoding cytolytic molecules such as *GZMB*, *PRF1* and *NKG7* and showed enrichment for cytotoxicity gene signatures (Fig. 1F, Fig. 2B and C; Additional file 1: Fig. S1F; Additional file 2: Tables S3 and S4) [4], suggesting that clusters 2 and 4 are cytotoxic clusters. The presence of mixture of cells expressing *CD4* and *CD8* transcripts in both the cytotoxic clusters 2 and 4 suggest possible similarities between CD4-CTLs and CD8-CTLs.

**Fig. 2.**
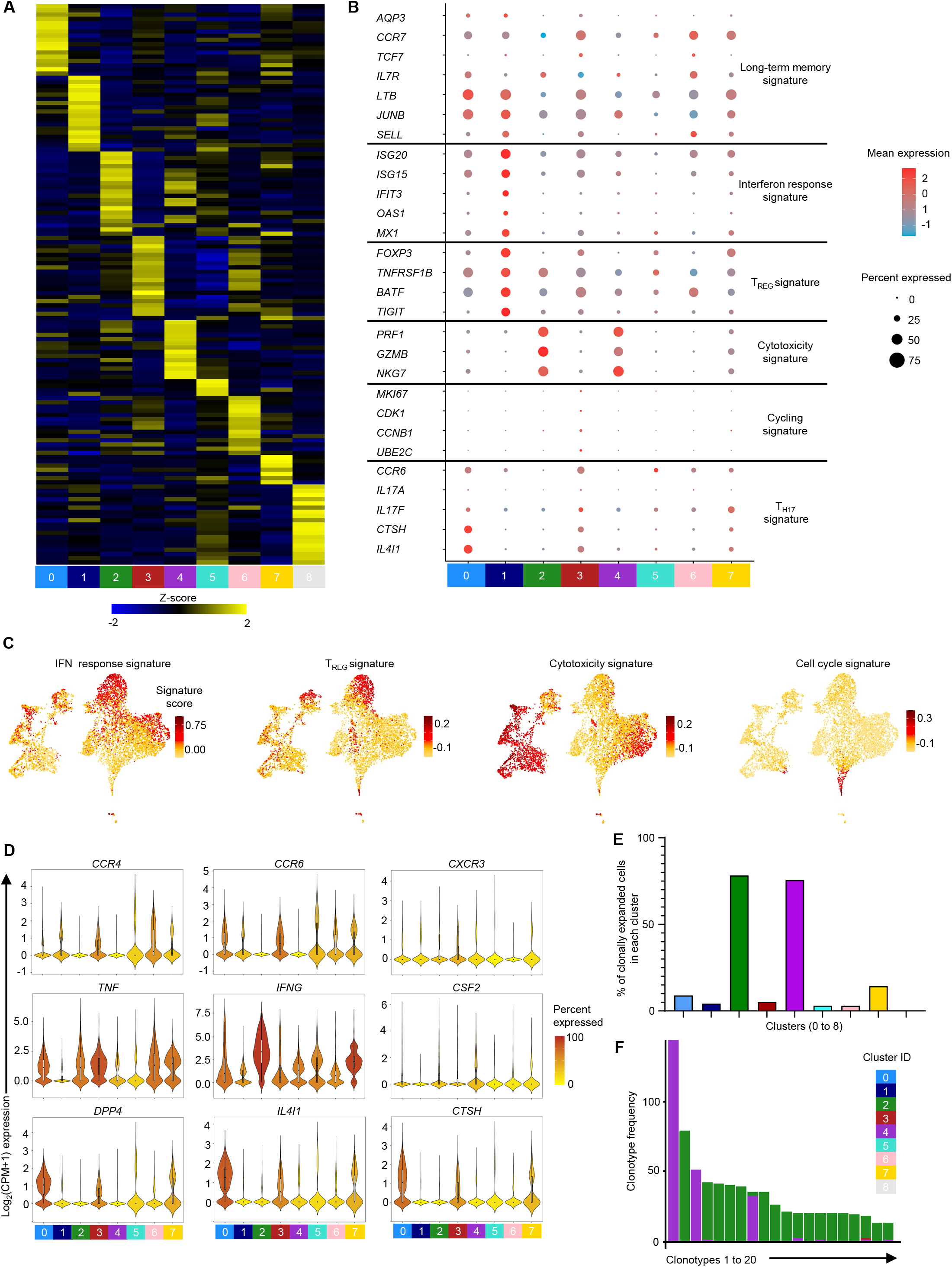
hCMV-reactive T cell subtypes and their TCR clonal diversity. (A) Heatmap shows the expression of the most significantly enriched transcripts in each cluster. The top 200 transcripts are shown based on adjusted *P* value < 0.05, log_2_ fold change > 0.25 and > 10% difference in the percentage of cells expressing selected transcript between two groups of cells compared. Each column represents the average expression of all cells for a given cluster. Colour scale represents z-score for the expression of each transcript across clusters. (B) Dot plots show the mean expression (colour) and percentage of expressing cells (size) for selected marker gene transcript in each cluster. (C) 2D UMAP plots show the IFN response, T_REG_, cytotoxicity and cell cycle signature score for each cell. (D) Violin plots show the normalized expression level (log_2_(CPM+1)) of the indicated differentially expressed transcript. Colour scale shows percentage of cells expressing the indicated transcript for each cluster. (E) Bar graph shows the percentage of cells with expanded clonotypes (frequency≥2) across 9 clusters. (F) Bar graph shows the frequency of 25 most expanded clonotypes across different clusters. For B and D, clusters with <1% of the total cells are not shown (cluster 8=56 cells).

We next interrogated the single-cell transcriptome of hCMV-specific T cells for functional T helper (T_H_) cell subtypes such as T_H_1, T_H_17 and T_H_1/17, that are known to have role in viral infections [3, 44]. Multiple clusters showed expression of T_H_ subset-specific transcripts: T_H_17 (*CCR4*, *CCR6*, *IL4I1*, *CTSH*), T_H_1 (*CXCR3*, *TNF*, *IFNG*, *CSF2*) and T_H_1/17 (*CXCR3, DPP4*) [9, 10] (Fig. 2D; Additional file 2: Table S4). Interestingly, we found several clusters co-expressing transcripts from different T_H_ subsets, suggesting each cluster may have cells belonging to more than one T_H_ subtype. Independently, module enrichment analysis using T_H_ subset-specific gene sets corroborated these results (Additional file 1: Fig. S2A; Additional file 2: Table S3). The T_H_1-specific gene sets were enriched in cluster 2 and 4, the cytotoxic clusters (Additional file 2: Fig. S2A). Considering there are several overlapping genes between T_H_1 and cytotoxic gene sets, this observation is in alignment with the known literature [4, 9, 10]. T_H_17 signature genes showed highest enrichment in cluster 0, followed by clusters 7, 3 and 1. T_H_1/17 signatures were mostly enriched in cluster 0 followed by clusters 5 and 7 (Additional file 1: Fig. S2A) [9, 10]. Together these observations indicate that each hCMV-specific T_H_ subset is at different developmental stages of the memory (long-term memory and effector memory) and clustering was predominantly influenced by the developmental stages of memory T cells

### TCR clonally expanded hCMV-specific T cells are majorly observed in cytotoxic clusters

To understand the TCR clonal diversity amongst the hCMV-specific T cells we analysed the single-cell TCR-Seq data generated from the same cells. We recovered TCRs from 83.56% (5765 cells of 6899 cells) of the total QC-ed cells in our data set, of which 21.40% (1234 of 5765 cells) were clonally expanded (Fig. 2E; Additional file 2: Table S5). Several clusters shared TCR clonotypes, indicating a shared developmental lineage between these clusters (Additional file 1: Fig. S2B). A maximum of 14 clonotypes were shared between clusters 2 and 4, followed by clusters 0 and 3 sharing 10 clonotypes (Additional file 1: Fig. S2B). Cells in cluster 2 and 4 showed highest clonal expansion with ∼78% and ∼76% of the cells being expanded respectively (Fig. 2E; Additional file 2: Table S5). As high as 76% of the total expanded cells (938 of 1234) were found in cluster 2 (50%) and cluster 4 (∼26%) (Additional file 2: Table S5). A few top expanded clonotypes were found in as many as 144, 79 and 51 cells from clusters 2 and 4, indicating the preferential expansion of a few specific clonotypes (Fig. 2F; Additional file 1: Fig. S2C). Interestingly, close to 58% of the expanded cells carried only 20 clonotypes (Fig. 2F). Both CD4-CTLs and CD8-CTLs (T_EMRA_) are shown to be clonally expanded in several diseases including CMV in both humans and animal models [2–4, 45, 46]. Clonal expansion and enrichment for cytotoxic signature in clusters 2 and 4 emphasizes the role for cytotoxic cells in hCMV-specific protective immune response [16–20].

### Single-cell transcriptome analysis of hCMV-specific T_REG_ cells

T_REGs_ are characterised by the higher expression of CD25 (IL2RA) and the transcription factor FOXP3, but a lower expression of IL7Rα (CD127) in humans [47]. CMV-specific T_REGs_ have been reported in both mouse models and humans following primary infections and are shown to produce IL10, a suppressive cytokine [2, 6, 36–39, 48–50]. However, the detailed molecular signatures of hCMV-specific memory T_REGs_ have not been well described. The transcript encoding the T_REG_ specific transcription factor FOXP3 was differentially expressed in clusters 1 (85% of cells) and 7 (Fig. 2B, Fig. 3A and B; Additional file 2: Table S4), however though not differentially expressed, a portion of cells from cluster 5 showed higher *FOXP3* expression (41% of cells), while various other clusters (clusters 0 and 3) showed lower levels of *FOXP3* expression (Fig. 3A and B). Considering FOXP3 can be transiently expressed on TCR activated cells, we examined the co-expression of other T_REG_ related genes and *FOXP3*. The cells co-expressing *FOXP3* - *IL2RA*, and *FOXP3* - *IKZF2* were enriched in cluster 1 and portion of cluster 5, and as expected they showed lower expression of IL7Rα (Fig. 1C and Fig. 3A) [47]. Clusters 1 and 5 were also characterized by the higher expression of other T_REG_-specific transcripts such as *TIGIT*, *BATF*, *TNFRSF1B* and the Ikaros Zinc Finger (IkZF) transcription factor (TF) family members *IKZF1* encoding Ikaros, *IKZF2* encoding Helios, *IKZF3* encoding Aiolos, *IKZF4* encoding Eos (Fig. 2B and Fig. 3C). The overall T_REG_ gene signature was enriched in cluster 1 and fraction of cluster 5 (Fig. 2C). Together these results show that T_REG_ cells are predominantly in cluster 1 and portion of cluster 5 and the cells from clusters 0 and 3 are potentially non-T_REG_ cells, transiently expressing low levels of *FOXP3* in response to TCR activation. We next interrogated if the hCMV antigen-specific T_REGs_ in our data set were indeed memory T_REGs_. Based on the expression of CD45RA, the FOXP3^+^ T_REGs_ have been classified as resting or naïve T_REGs_ (FOXP3^lo^CD45RA^+^) and effector T_REGs_ (FOXP3^hi^CD45RA^-^) [51, 52]. Further the effector T_REGs_ have been classified as central and effector memory cells based on the expression of CCR7, CD27 and CD28 [53]. The T_REGs_ in our data set were characterized by the lack of expression of CD45RA, and a moderate expression of CCR7, *CD27* and *CD28*, indicating that they are mostly central and effector memory T_REGs_ rather than naïve T_REGs_ (Fig. 1C and D).

**Fig. 3.**
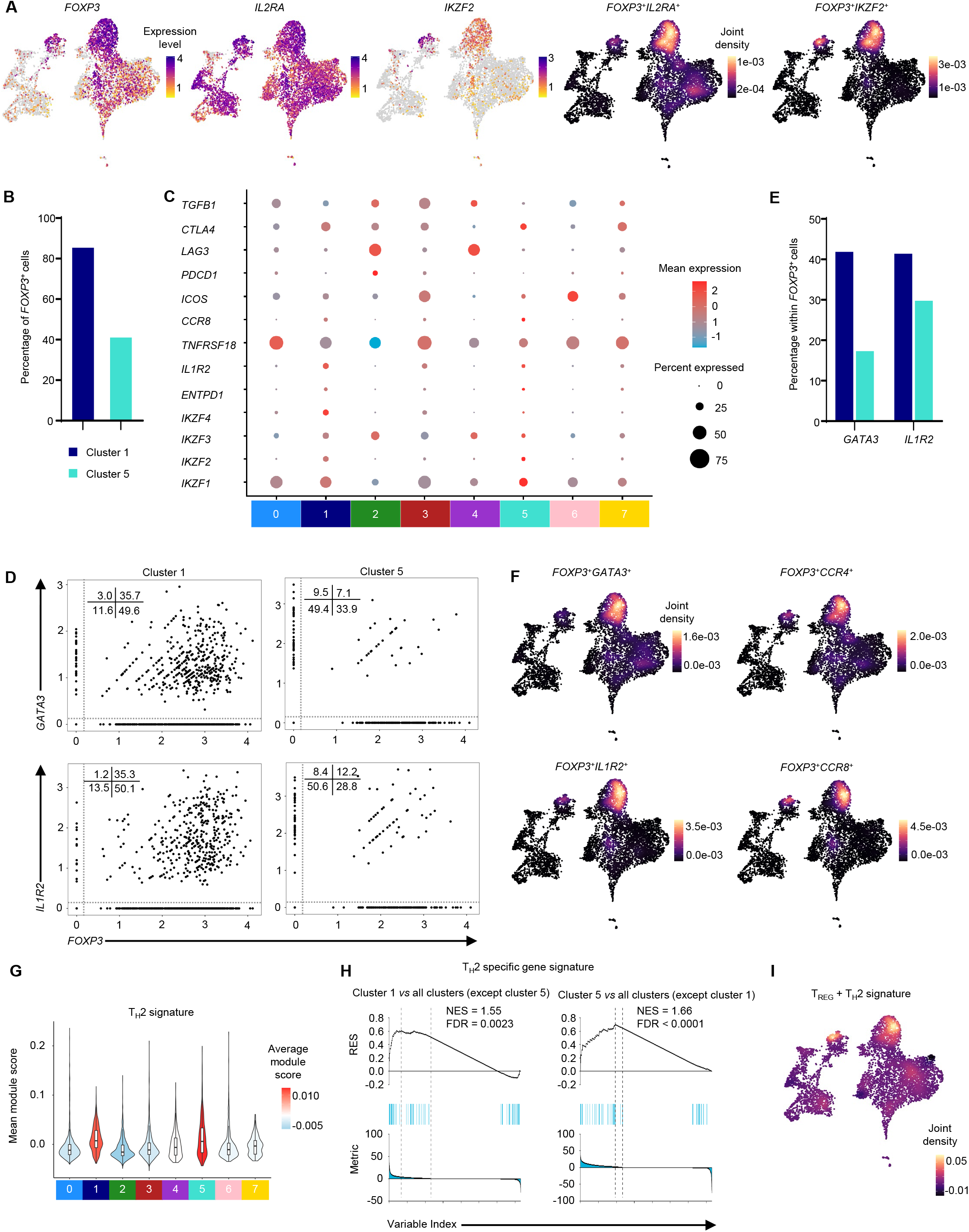
Single-cell transcriptomic analysis of hCMV-reactive T_REGs_. (A) 2D UMAP plots of single-cell RNA-Seq data show the expression of the *FOXP3*, *IL2RA*, *IKZF2* (left) and joint expression of *FOXP3* with *IL2RA* and *FOXP3* with *IKZF2* (right 2 plots) transcripts. Scale represents log_2_ normalized (CPM+1) counts for expression values and scaled density for joint expression plots. Cells with zero counts for the indicated transcripts are shown in grey colour. (B) Bar graph shows the percentage of cells expressing *FOXP3* transcript (log_2_(CPM+1)>0) in clusters 1 and 5. (C) Dot plots show the mean expression (colour) and percentage of expressing cells (size) for selected marker gene transcripts in indicated clusters. Clusters with <1% of the total cells are not shown (cluster 8=56 cells). (D) Scatter plots showing normalized co-expression level (log2(CPM+1)) between *FOXP3* and *GATA3* and *FOXP3* and *IL2RA* transcripts in the indicated clusters. (E) Bar graph shows the percentage of cells expressing *GATA3* or *IL1R2* transcripts (log_2_(CPM+1)>0) within *FOXP3* transcript expressing cells in clusters 1 and 5. (F) 2D UMAP plots show joint expression of *FOXP3 + GATA3*, *FOXP3 + CCR4* and *FOXP3* + *IL1R2*, *FOXP3 + CCR8*. (G) Violin plot shows the module enrichment score for T_H_2-specific gene set across 8 clusters. Colour scale indicates mean module score. Clusters with <1% of the total cells are not shown (cluster 8=56 cells). (H) Gene set enrichment plot for T_H_2 specific gene sets in cluster 1 and cluster 5 versus all the other cluster. RES: Running Enrichment Score; NES: Normalized Enrichment Score; FDR: False Discovery Rate. (I) 2D UMAP plot shows joint expression of T_H_2-specific gene set and allergy induced T_REG_-specific gene set. Scale represents density for joint expression plot.

Although, T_REGs_ expressing FOXP3, are known for their suppressive function, many reports have shown the presence of T cells expressing FOXP3 transiently, in response to TCR activation that are non-suppressive in nature. Hence, we assessed if the hCMV-specific T_REGs_ in our data set are expressing markers associated with the suppressive nature of T_REGs_. The molecular signatures of T_REGs_ found in clusters 1 and 5 indicated their potential suppressive nature, compared to cells expressing low levels of *FOXP3* observed in clusters 0 and 3 (Fig. 3A). The T_REGs_ in clusters 1 and 5 were *CD137*^+^*CD154^lo^*, the activation induced characteristic of T_REGs_ implicative of epigenetically stable antigen-activated T_REGs_ that retain the high suppressive potential in humans after *in vitro* or *ex vivo* activation (Fig. 1D and G) [54]. The T_REGs_ in clusters 1 and 5, showed higher expression of *TIGIT*, *TNFRSF18* (GITR) and *CTLA-4*, genes that are associated with T_REGs_ containing high affinity TCRs, further suggesting their antigen-specificity (Fig. 2B and Fig. 3C). The T_REGs_ in clusters 1 and 5 also showed higher expression of *ENTPD1* (CD39), an ectoenzyme ATP apyrase, *CCR4*, *CTLA4*, Helios (*IKZF2*) and FAS (CD95), molecules that are reported to be associated with effector and suppressive function (Fig. 1C, Fig. 2D and Fig. 3C) [55–57]. A suppressive function of Helios in T_REGs_ has been suggested, where Helios^+^FOXP3^+^ T_REG_ cells produced lower amounts of T_H_17-specific cytokine, IL17, compared to Helios^-^FOXP3^+^ T_REG_ cells, hence promoting the formation of T_REGs_ rather than T_H_17 subtype [57]. Consistent with these observations, we found that clusters 0 and 3 that expressed *IL17F* and *IL17A*, expressed low levels of *FOXP3*, but lack the expression of *IKZF2* (Helios) and these cells showed higher expression of *CD154* and lower expression of *CD137* (Fig. 2B, Fig. 3A and C) [54, 57]. Together these observations suggest that the low *FOXP3* expressing cells in clusters 0 and 3 are potentially the induced/transient T_REGs_ that are possibly expressing FOXP3 transiently in response to activation and are non-suppressive in nature. Conversely, T_REG_ cells in cluster 1 and 5 are *Helios^+^FOXP3^+^*, and did not express *IL17A* and *IL17F*, indicative of suppressive nature (Fig. 2B, Fig. 3A and C). The lack of expression of exhaustion markers *PDCD1* and *LAG3* further suggests these are not associated with anergy or exhaustion (Fig. 3C). Together these results provide multiple evidences to suggest that the T_REGs_ in clusters 1 and 5 are effector memory T_REGs_ and are potentially suppressive in nature. One of the mechanisms through which T_REGs_ execute their suppressive function on effector T cells is through cytokines such as IL10, TGFβ and IL35 (reviewed in [58]). The T_REGs_ in cluster 1 lacked or showed low expression of these suppressor cytokine transcripts *TGFB1*, *IL10* and *IL35*, indicating that cells in cluster 1 may not be actively suppressing the hCMV-specific effector T cells, at least in the experimental setting used in the study (Fig. 3C; Additional file 2: Table S4).

We next interrogated if the T_REG_ subsets in our data set, that showed enrichment for molecules linked to their suppressive function, are not suppressive towards hCMV-specific effector T cells, then which cell types are they targeting in hCMV-specific immunity. Studies using mouse models have shown that T_REGs_ can show remarkable functional plasticity and execute their suppressive function by expressing the TF and the chemokine receptor of the target T_H_ subtypes [14, 59–62]. For example, the T_REGs_ suppressive towards T_H_1 will express T-bet (encoded by *TBX21*) and CXCR3 (T_H_1-T_REGs_), while the ones toward T_H_2 will express GATA3 and CCR4 (T_H_2-T_REGs_), albeit at a reduced level compared to the T_H_ subtype itself, without necessarily themselves making the T_H_ subtype-specific cytokines [14, 60–62]. Interestingly the differential gene expression analysis revealed that the T_REGs_ in cluster 1 and 5, are characterized by the higher expression of T_H_2-specific TF *GATA3* and chemokine receptor *CCR4* along with *FOXP3* (Fig. 3D-F). GATA3 has been shown to be important for FOXP3 expression and T_REG_ functions [61]. Hence, we independently examined the overall T_H_2 subset-specific gene signatures using module enrichment score and gene set enrichment analysis (GSEA). We found that both cluster 1 and 5 are enriched for T_H_2 subset-specific gene signatures (Fig. 3G and H), suggesting the T_REGs_ in cluster 1 and 5 are most probably T_H_2-like T_REGs_. A recent study that analysed the T cells in pan cancers has also shown the presence of T_H_2-like T_REGs_ that were marked by a higher expression of *GATA3* [63]. Clusters 1 and 5 were also characterised by the enrichment of IFN response signature genes (T_REG-IFN_) (Fig. 2B and C). In asthma and house dust mite allergy, both T_H_ and T_REG_ subsets enriched for IFN response gene sets showed dampened T_H_2 response in healthy individuals with no prior history of allergy or asthma [5]. Interestingly, in our data set, a joint enrichment analysis using the house dust mite allergy-specific T_REG_ gene signature and T_H_2-specific gene signature showed enrichment in cluster 1 and 5 (Fig. 3I) [5]. Together these results suggest that T_REGs_ in clusters 1 and 5 are likely T_H_2-T_REGs_ with IFN response signatures that are potentially suppressive towards T_H_2. The T follicular regulatory cells (T_FRs_) suppress T_FH_ responses and are characterized by the expression of IL1R2 and CCR8 [3, 64, 65]. The hCMV-reactive T_REG_ cells in clusters 1 and 5 co-expressed *IL1R2* and *CCR8* transcripts along with *FOXP3* transcripts, hence, may represent T_FR_ cells (Fig. 3D-F) [3, 64, 65]. These interesting observations warrant further investigation to understand the relevance of T_H_2-T_REGs_ and T_FRs_ in CMV immunity, especially when hCMV-reactive conventional (non-T_REGs_) T_H_2 and T_FH_ cells were absent in our data set.

### hCMV-reactive CD4-CTLs and CD8-CTLs show similar transcriptomic profile

Majority of the cells in clusters 2 and 4 and fraction of cells in clusters 0 and 7 were characterized by the high expression of transcripts encoding cytolytic molecules such as Granzyme B (*GZMB*), Perforin (*PRF1*), Granulysin (*GNLY*), and transcription factors *HOPX* and *ZEB2* and showed enrichment for cytotoxicity and T_EMRA_-specific signature genes (Fig. 1F, Fig. 2B and C, Fig. 4A and B) [3, 4, 16–20]. Interestingly, these cytotoxic clusters could not be differentiated based on the expression of *CD4* or *CD8A* and *CD8B* transcripts, but by other cytotoxicity related molecules (Fig. 1E and Fig. 4A-C). The cytolytic granules have been shown to contain chemokines such as CCL3, CCL4 and CCL5 in addition to the classical mediators of cytotoxicity (GZMB, PRF1, GNLY), in both viral and bacterial infections, indicating their role in cytolysis [66, 67]. Cells in cluster 2 showed higher expression of the transcripts encoding the cytokines such as *TNF* and *IFNG*, chemokines such as *CCL3*, *CCL4*, *CCL4L2*, *XCL1*, *XCL2*, while cluster 4 showed higher expression of *CCL5* (Fig. 4B and C). The chemokine receptor *CCR5*, the receptor for chemokines *CCL3*, *CCL4* and *CCL5*, and *IFNGR1*, encoding the receptor for IFNG, was expressed by the cytotoxic cells in cluster 0, suggesting these cells may be responding to the cytokines and chemokines made by the cells in cytotoxic clusters 2 and 4 and are potentially the intermediate population (Fig. 4A and B; Additional file2: Table S4) [68]. This is further supported by the single-cell trajectory analysis, where on the pseudo time scale cluster 0 appears before clusters 2 and 4 (Fig. 4D). These results suggest that the cytotoxic cells in cluster 0 are potentially an intermediate population in the CTL lineage and these cells are mostly T_EM_ cells as they mostly lack the expression of CD45RA and CCR7 (Fig. 1C).

**Fig. 4.**
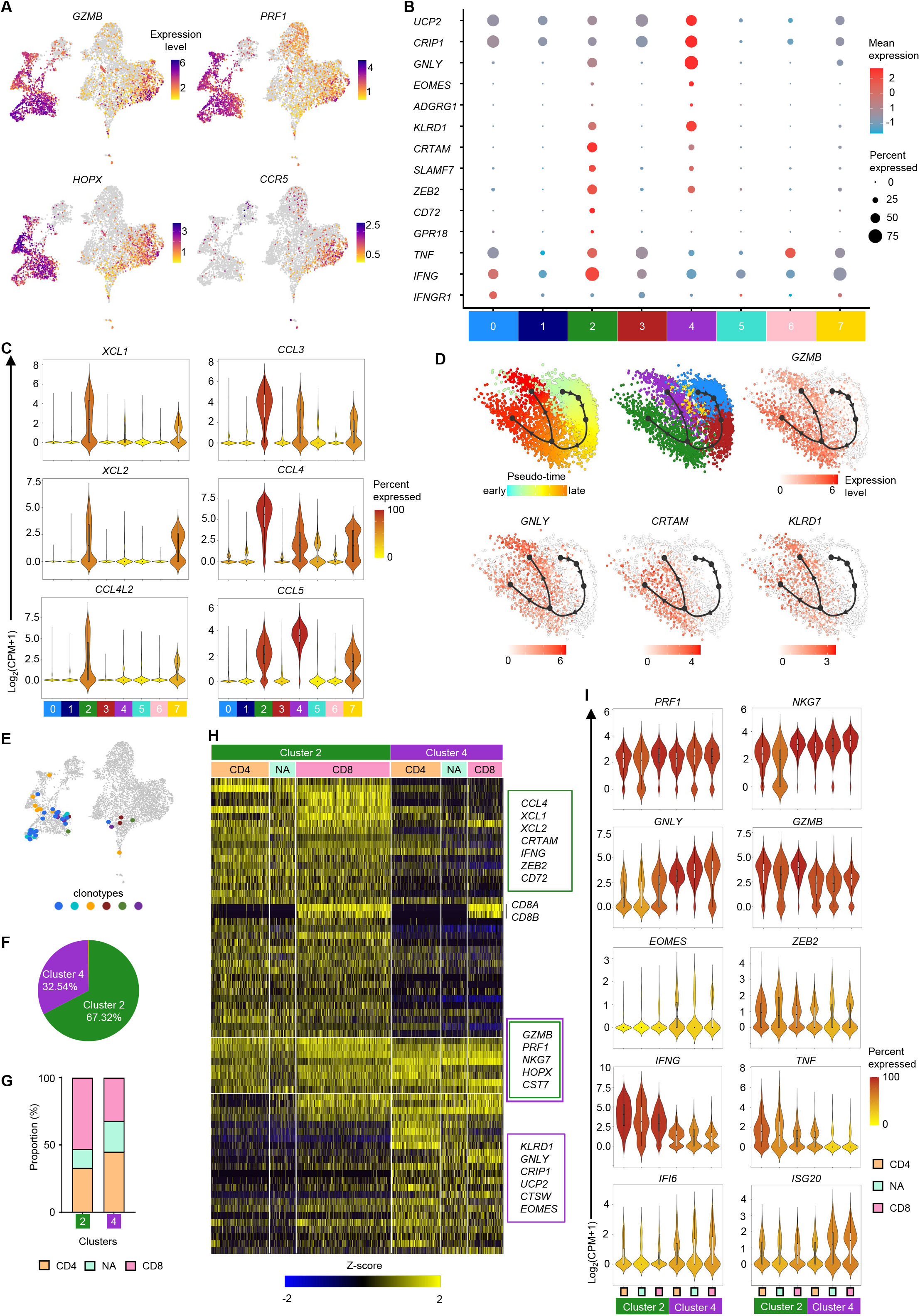
Single-cell transcriptomic analysis of hCMV-reactive CD4-CTLs and CD8-CTLs. (A) 2D UMAP plots of single-cell RNA-Seq data show the expression of the indicated transcripts. All expression counts for transcripts are log_2_ normalized (CPM+1). Cells with zero counts for the indicated transcripts are shown in grey colour. (B) Dot plots show the mean expression (colour) and percentage of expressing cells (size) for selected marker gene transcripts in each cluster. (C) Violin plots showing the normalized expression level (log_2_(CPM+1)) of the indicated differentially expressed transcripts across 8 clusters. Colour scale shows percentage of cells expressing the indicated transcript for a given cluster. (D) Trajectory analysis using slingshot shows the distribution of cells from indicated clusters (middle) on a pseudo-scale (left), and expression levels (scale) of the *GZMB*, *GNLY*, *CRTAM* and *KLRD1* transcripts. (E) 2D UMAP plot shows the cells with same clonotypes (colours) shared between cluster 3 and, clusters 2 and 4. (F) Pie chart shows the distribution of percentage of cells expressing the top 20 expanded clonotypes in clusters 2 and 4. (G) Stacked bar graph shows the proportion of CD4, CD8 and NA (not assigned) T cells in each indicated cluster (refer to methods section for classification of cells). (H) Heatmap shows the normalized expression of 50 differentially expressed transcripts from each clusters 2 and 4 when compared to the rest of the clusters. (I) Violin plots show the normalized expression level (log_2_(CPM+1)) of the indicated differentially expressed transcript across indicated T cell subsets in clusters 2 and 4. Colour scale shows percentage of cells expressing the indicated transcript. For B and C, clusters with <1% of the total cells are not shown (cluster 8=56 cells).

Compared to cluster 4, cluster 2 cells showed higher expression of *SLAMF7*, *ZEB2*, *GZMB*, *GPR18*, *CD72*, *PDCD1* (PD1), *CRTAM* that are known to be expressed in terminally differentiated effector cells in viral infections and various cancers [3, 69–72] (Fig. 2B, Fig. 3C, Fig. 4B and D). On the contrary, cluster 4 was characterized by the higher expression of *GNLY*, *EOMES*, *ADGRG1* (GPR56), *KLRD1*, *UCP2* and *CRIP1* (Fig. 2B, Fig. 4B and D), that are mostly not associated with terminally differentiated effector T cells. For example, CRIP1 has been shown to be anti-apoptotic in cancers and the antigen-specific cells expressing UCP2 survive better in HIV infection through metabolic reprogramming [73, 74], suggesting the possible role of these genes in providing survival advantage to cells in cluster 4. A coordinated expression of CCL5 followed by GNLY and PRF1 provided host-defence against *Mycobacterium tuberculosis* (Mtb) infection, where the expression of CCL5 attracted the Mtb infected macrophages that in-turn triggered the expression of GNLY and PRF1 in CD8 T cells [75]. Further, a delayed onset of cells expressing GNLY has been shown in multiple infections [76, 77]. These results suggest that the cytotoxic cells could be at different stages of their development or effector function. The differential expression of many of these cytolytic molecules between clusters 2 and 4, imply that the cells in cluster 2 are potentially the terminal effector cells while those in cluster 4 are pre-effector cells. This observation is further strengthened by the TCR analysis (Fig. 2E and F, Fig. 4E and F; Additional file 1: Fig. S2B and C). The 20 most expanded clonotypes were found majorly in cluster 2 and 4 (Fig. 2F; Additional file 1: Fig. S2C). Cluster 2 is relatively more clonally expanded than cluster 4, where over 67% of the cells harbouring the 20 most expanded clonotypes were found in cluster 2 (Fig. 4F). The trajectory analysis also showed that on a pseudo-time scale, cluster 2 and 4 appears to bifurcate from a common origin of cluster 0 (Fig. 4D). Further, the pathway enrichment analysis of genes differentially expressed between clusters 2 and 4 revealed an enrichment for cytokine and chemokine response pathways in cluster 4, while cluster 2 showed enrichment of pathways associated with pathogen induced cytokine storm (Additional file 1: Fig. S3A and B). The effector T cells utilize glycolysis, as the preferred metabolic pathway that play an important role for cytokine secretion and lytic nature of these effector cells [78–80]. The cells in cluster 2 showed enrichment for many metabolic pathways associated with effector cells such as glycolysis and HIF1α signalling as well as pathways related to anti-viral responses (Additional file 1: Fig. S3A). Together, these combined analysis of the transcriptome, TCR clonotype expansion and sharing along with the trajectory analysis provides the evidence to support the idea that cluster 4 cells are pre-effectors while cluster 2 cells are effectors.

Next to assess if the observed differences between the CTL clusters 2 and 4 are influenced by the proportion and the characteristics of CD4-CTLs and CD8-CTLs, we compared the expression of these transcripts between CD4 and CD8 T cells within clusters 2 and 4. The proportions of CD4 and CD8 T cells within clusters 2 and 4 were comparable (Fig. 4G), indicating that the disproportionate distribution of CD4^+^ and CD8^+^ T cells is not the cause of the observed differences between clusters 2 and 4. Overall, the transcriptome profile of CD4, CD8 or cells that could not be assigned as CD4 or CD8 T cells (NA) within each cluster did not vary except for *CD8A* and *CD8B* transcripts (Fig. 4H and I). Transcripts such as *PRF1* and *NKG7* that showed comparable expression between cluster 2 and 4, followed the same pattern even when these two clusters were further sub-divided based on the CD4 and CD8 T cell sub-type. The transcripts that showed higher expression in cluster 2 (*XCL1*, *ZEB2*, *CRTAM* and *GZMB*) and cluster 4 (*CCL5*, *GNLY*, *EOMES*), followed similar pattern of expression in CD4 and CD8 T cells within each of these effector and pre-effector clusters (Fig. 4H and I). Both CD4 and CD8 T cells from cluster 4 showed higher expression of IFN response genes *ISG20* and *IFI6* while those in cluster 2 showed higher expression of cytokines *TNF* and *IFNG* (Fig. 4H and I). We observed that the cells in clusters 2 and 4 have distinct gene expression pattern that is not influenced by either CD4 or CD8 T cell proportion (Fig. 4H and I). Overall, our data has revealed that both hCMV-reactive CD4-CTLs and CD8-CTLs share similar transcriptome profile and can be classified as effectors and pre-effectors of CTLs.

## Discussion

A significant world population carries the latent hCMV infection, without showing any symptoms or causing the disease. However, latent infection poses a huge risk of reactivation in immune-compromised individuals such as those undergoing transplant or infants born to hCMV seropositive mothers. Although there is very little risk of the disease occurrence in immune-competent individuals, the latent CMV infection puts a huge burden on the immune system, especially for the elderly, where significant proportions of the T cells are found to be hCMV-specific [46]. Identifying and understanding the characteristics of the hCMV-specific T cell immune memory components in individuals carrying the virus in latent form, representing protective immune response, can be of high value to design therapy or vaccines to control CMV infection upon reactivation or maternal to fetal transfer.

Our understanding of *ex vivo*, antigen-specific human T cell responses to pathogens such as hCMV remains limited, due to the rarity of the cells, the complexity of the biology, and technical challenges of identifying overall antigen-specific CD4^+^ memory T cells, since they can be extremely heterogeneous in the expression of activation associated markers and cytokines. An added layer of complexity comes from HLA-restriction of individuals and the prior knowledge about the immune-dominant peptide, limiting the use of pMHC multimers. Most studies have focused on isolating the total CD4^+^ or CD8^+^ T cells reactive to hCMV, without considering the differences in memory proportions that can inadvertently bias the analysis towards analysing the predominant memory compartment within CD4^+^ T cells (T_CM_ or T_EM_) and CD8^+^ T cells (T_EMRA_). Our strategy to isolate antigen-specific T cells based on the combination of TCR activation dependent markers (CD154 and CD137) across different memory subsets followed by high resolution single-cell RNA-Seq paired with single-cell TCR analysis allowed us to interrogate and compare a larger pool of hCMV (pp65 and IE-1)-reactive cells from different memory subsets across 10 seropositive and peptide-responsive donors. This unbiased approach enabled us to overlay the heterogeneity in T cell memory subsets classified based on developmental stages and functional subsets along with their TCR dependent activation markers. The expression of CD154 and CD137 in different set of cells that further differentially co-expressed one or more T cell activation associated markers such as CD69, CD25, OX40, warrants for careful selection of activation markers while analysing antigen-specific cells *ex vivo*. In this study, we observed that though activated (based on the expression of *CD154*, *CD137*, *CD69*, *CD25*, *OX40*), most cells, with the exception of effector cells, did not actively make cytokines and those that made also showed heterogeneity for the cytokine profile. Many studies previously have also relied on cytokine profiles to analyse *ex vivo* isolated antigen-specific cells and hence have mostly analysed effector memory T cells.

The single-cell RNA-Seq of a large number of hCMV reactive cells isolated from 10 different donors, identified several interesting functional subsets; T_H_17, T_REGs_, CD4-CTLs and CD8-CTLs. Of particular interest were CD4- and CD8-CTLs, that clustered together. The two cytotoxic clusters observed could not be distinguished based on the CD4- and CD8-T cell specific transcripts *CD4* or *CD8A*/*CD8B*, but on the basis of other cytotoxic molecules that were linked to different states of effector functions [72]. The cluster that was predominantly making cytokine and chemokine (*IFNG*, *TNF*, *XCL1*, *XCL2*, *CCL3*, *CCL4*) also expressed transcription factor (TF) *ZEB2*, the TF known to be associated with late effector function [4, 72]. The cytotoxic cells in both clusters 4 and 0 appeared to respond to the cytokines and chemokines by expressing their receptors (*IFNGR1*, *CCR5*). However, the cytotoxic cells across these clusters expressed molecules linked to cytolytic granules (*PRF1*, *GZMB*, *GNLY*), and transcription factor *HOPX*. The differential expression of many of the molecules linked to cytotoxicity and effector functions, suggests the effector and pre-effector like subsets. These subsets, possibly at different developmental states, might exist to ensure a continuity in immune response during the entire course of the infection.

Another interesting discovery in our data set is the identification of T_REGs_ with suppressive ability that showed enrichment for the IFN response gene signatures (T_REG-IFN_) which has been shown to be associated with a dampened T_H_2 response in allergy [5]. The enrichment of T_H_2 signatures as well as the expression of *GATA3* and *CCR4*, the T_H_2 associated transcription factor and chemokine receptor by the T_REGs_ further suggested that these could be suppressive towards the T_H_2 subset. Although a very recent study in pan cancer has also identified this subset, to the best of our knowledge, presence of T_REGs_ enriched for IFN response genes has not been reported earlier in viral infections in humans [63]. Based on these observations, we hypothesize that during the initial stages of the infection where the expansion of effector cells is necessary, T_REGs_ may not be executing their suppressive function towards these effectors, instead, suppressing other subtypes of no relevance to the disease. However, further kinetic studies in both animal models and longitudinal human cohorts of different types of diseases such as viral and bacterial infections, allergy and cancers are necessary to verify or refute this hypothesis.

## Conclusion

In conclusion, using the single-cell multi-omics approach (transcriptome, TCR repertoire and expression of a few selected proteins), our study has identified an unprecedented amount of heterogeneity in hCMV-specific CD4^+^ memory T cell subsets. The study identified novel subsets such as T_REG-IFN_ and pre-effectors and effectors of both CD4-CTLs and CD8-CTLs. The knowledge base can serve as an important resource for designing vaccines and therapeutic strategies to control CMV infections, especially in immune-compromised individuals and infants of seropositive mothers.

## Materials and Methods

### Study subjects

Study subjects were healthy adult donors who donated blood at the Safdarjung Blood Bank during 2019-20. The donors consented for use of the plasma and buffy coat samples for research. Donors were HIV-negative and had no history of Hepatitis C infection. The median age was 30.5 (24-39) and all these donors were male. Approval for the use of this material for research was obtained from the institutional human ethics committees from both National Institute of Immunology, New Delhi and Vardhman Mahavir Medical College (VMMC) and Safdarjung Hospital, New Delhi, India.

### PBMC processing

Peripheral Blood Mononuclear Cells (PBMCs) were isolated from buffy coat samples from healthy blood donors obtained from Safdarjung hospital blood bank by density gradient centrifugation using Ficoll-Paque Premium (GE Healthcare Biosciences). PBMCs were cryopreserved in 90% Fetal Bovine Serum (FBS) supplemented with 10% di-methyl sulfoxide (DMSO).

### CMV IgG analysis

ELISA (Enzyme Linked Immunosorbent Assay) was performed on the plasma samples from the donors obtained from the blood bank using Human Anti-Cytomegalovirus IgG ELISA kit (abcam, #ab108724) according to the manufacturer’s protocol to check for the donors’ sero-status. Donors with the absorbance values at least 10% higher than that of the cut-off control supplied in the kit, were considered positive. Based on this criteria we found 124 out of 126 donors in our cohort were hCMV IgG+; 98.4% positivity) (Additional file 1: Fig. S1A).

### PBMC stimulation

To analyse or isolate antigen-specific T cells, the ARTE assay was employed with few minor modifications [40, 41]. The hCMV-specific PepTivators (Miltenyi Biotec) derived from pp65 (UL83) and IE-1 (UL123) were used to stimulate PBMCs from CMV-seropositive donors, according to the manufacturer’s protocol. Briefly, PBMCs were thawed and treated with 50 U/mL Benzonase (Sigma) in RPMI before resting overnight in serum-free TexMACS medium (Miltenyi Biotec) supplemented with 1% Penicillin/Streptomycin at 37°C in a 96-well plate. Cells were stimulated for 6 or 24hrs at a density of 1 million/100 µL/well of 96 well plate, by the addition of hCMV peptide pools (pp65 and IE-1 PepTivators) at a final concentration of 1 µg/mL in the presence of a blocking CD40 antibody (1 µg/mL; Miltenyi Biotec) and CD28 antibody (1 µg/mL; eBioscience) [1, 40, 41]. Stimulation with αCD3/CD28 DynaBeads Human T cell activator (Invitrogen) (cell to bead ratio of 1:1) served as positive control while with water (vehicle control) served as unstimulated negative control. The stimulated cells were washed and stained with cocktail of fluorescent antibodies (Table S2A) and analysed by flow cytometry or processed for single-cell RNA-sequencing using 10X genomics as follows. The hCMV-peptide pool stimulated cells from each donor were washed and individually stained with Cell-hashtag (HTO:Hash Tag Oligo) TotalSeq C antibody (0.5 µg/condition; Biolegend, Table S2B) as per manufacturer’s recommendations for 30 minutes on ice, and washed twice with MACS buffer (PBS + 2% FBS + 2mM EDTA). HTO stained PBMCs from all 10 donors were then pooled and stained with oligo-tagged TotalSeq C antibody for cell surface proteins for T cell memory markers (CD45RA, CCR7, CD95 and IL7Rα) (0.5 µg/condition; Biolegend, Table S2B) for 30 minutes on ice. Following washing, the pooled cells were further stained with fluorescence-conjugated surface antibodies to enable FACS sorting of antigen-specific cells (Table S2A). Cells were directly FACS sorted without an intermediate MACS column enrichment step using FACSAria Fusion Cell Sorter (Becton Dickinson) in LoBind 1.5 mL microcentrifuge (Eppendorf) tubes with 1:1 FBS:PBS supplemented with recombinant RNase inhibitor (1:100, Takara). The live singlet gated CD3+ T cells were further gated as per the gating strategy shown in additional file 1: Fig. S1B. List of antibodies used are provided in additional file 2: Table S2. The flowcytometry data was analysed using FlowJo software (v10.8).

### Single-cell RNA-Seq

The sorted antigen-specific cells from various memory compartments were pooled in desired ratios and used for single-cell RNA-Seq and TCR-Seq assay using 10X Genomics kit, 5ʹ V(D)J with gene expression and cell surface protein expression v1.1 as per the manufacturer’s recommendations. Initial amplification of cDNA libraries and final libraries were performed for 15 cycles and 14 cycles respectively for gene expression library. V(D)J and cell surface protein libraries were generated corresponding to each 5ʹ gene expression library using 9 cycles of amplification. Libraries were sequenced on Illumina NovaSeq6000 sequencing platform for paired end (PE)150 reads.

### Single-cell RNA-Seq analysis

Sequenced reads from the 5ʹ single-cell gene expression and associated V(D)J library were mapped to the GRCh38 reference genome using 10X Genomics’ cellranger multi (v6.1.1). The cell surface protein quantification was run as part of the same pipeline. The Seurat package (v4.3) in R (v4.2.2) was used to perform all further downstream analyses [81].

For downstream analysis, only cells that passed the QCs were retained; cells with UMI (Unique Molecular Identifier) counts >500, number of genes >350 and percent mitochondrial genes <15% were considered as good quality cells. Further, cells were assigned as doublets, singlets or negatives based on the HTO counts using Demux with default parameters in Seurat package (v4.3) [82]. Singlets were assigned to their respective donor based on HTO count. Doublets were removed from downstream analysis. After applying all the quality control parameters, we retained 6899 cells out of 8062 cells for downstream analysis (85.57% cells passed the quality parameters). Cells from experiment 1 (3 donors) and experiment 2 (10 donors) were combined using the Seurat package after applying the QCs (Table S1).

For each cell, normalized UMI counts for each gene were obtained by dividing the raw counts by the total counts across all genes. This was multiplied by a scaling factor of 10,000 and log transformed to yield the final normalized values. Prior to clustering, we filtered genes by selecting for the top 2000 with highest variance across cells, using *FindVariableFeatures* with *selection.method*=”*vst*”. This data was then centered and scaled using *ScaleData*, and the top 50 principal components (PCs) were calculated. Top 18 PCs were selected using *ElbowPlot* and *JackStrawPlot* for downstream processing. This was used to build a shared nearest neighbour graph using *FindNeighbors*. The cells were clustered using *FindClusters*, with *algorithm=3* and *resolution=0.2*, determined using the *clustree* function. For visualization and clustering, a 2D UMAP projection of the top 18 PCs was obtained using the *RunUMAP* function, with *metric=”euclidean”* [81].

The cell surface protein tag data was normalized using the *NormalizeData* function in Seurat with *normalization.method*=”CLR”. Briefly, for each cell, the hashtag counts were divided by the geometric mean of counts of all unique hashtags prior to log transformation.

### Differential gene expression analysis

We identified differentially expressed genes using the MAST algorithm integrated within Seurat’s *FindAllMarkers* function using default parameters on log_2_(CPM+1) counts [81]. Genes in a cluster were reported as differentially expressed (DE) compared to every other cluster if their log_2_ fold change of expression was > 0.25 and BH-adjusted P-value was < 0.05.

### Gene module signature scores and Gene Set Enrichment Analysis (GSEA)

Gene module scores were computed using the *AddModuleScore* function in Seurat, with default parameters [81]. Briefly, for each cell, the module score is defined by the mean of the signature gene list (test set) after subtracting the mean expression of an aggregate of control gene lists. Control gene lists (same number as test gene list) are sampled from bins created based on the level of expression of the signature gene list. GSEA was performed using Qlucore Omics Explorer 3.8 software package [83]. List of the genes used are provided in supplementary table S3.

### Trajectory inference

Single cell trajectories were computed using the slingshot algorithm within the dynverse/dyno (v0.1.2) suite of trajectory inference pipelines [84]. Raw and normalized UMI counts from cells of clusters 0, 2, 3, 4 and 7 were combined using the dynverse’s wrap_expressions function. Cells were assigned groups by cluster and the number of expected start and end states were set to 1 using *add_prior_information*. Trajectory inference was run using *infer_trajectories* with default parameters, with method = ti_slingshot(). We did not specify any start or end cells for the trajectory.

### Pathway enrichment analysis

DE genes between clusters 2 and 4 were fed to Ingenuity Pathway Analysis software from Qiagen. Enrichment pathways were sourced from Ingenuity core enrichment pathways.

### TCR analysis

For each cell, all information about clonality, expressed TCR chains, their sequences and corresponding UMI counts were obtained as part of the standard 10X Genomics’ cellranger multi pipeline (Table S5).

### Assigning CD4 and CD8 T cells

For cells in cluster 2 and 4, we assigned cells as CD4^+^ T cells, or CD8^+^ T cells, first based on the *CD4* and *CD8A*/*CD8B* transcript counts. Cells with > 0 counts for *CD4*, were assigned as CD4^+^ T cell (333 cells). For CD8^+^ T cells, we considered the cell as CD8^+^ T cell (735 cells) if the sum of all three CD8 transcripts (*CD8A*, *CD8B*, *CD8B2*) was > 0 and *CD4* transcripts were undetectable. Cells (556) which could not be assigned as either CD4 or CD8 T cell based on this criterion, were further assigned as CD4^+^ (274) and CD8^+^ (12) T cells if they shared the clonotypes with an already assigned CD4^+^ or CD8^+^ T cell. Cells that could not be assigned to either CD4 or CD8 category based on either of these criteria, were called as NA (not assigned) cells (294). Based on these criteria, we had a total of 328 CD4^+^ T cells, 536 CD8^+^ T cells and 149 NA cells in cluster 2 (total 1013 cells) and 279 CD4^+^ T cells, 199 CD8^+^ T cells and 145 NA cells in cluster 4 (total 623 cells).

### Statistical tests

All the statistical tests were done using GraphPad Prism v9.0. Mann Whitney u-test was used to perform pairwise comparison and *P*-value <0.05 was considered significant.

### Data and Code availability

Scripts will be made available on GitHub (https://github.com/ImmunogenomicsLab-NII/CMV-HD). Sequencing data for this study is deposited onto the Gene Expression Omnibus with accession number GSE235604.

## Acknowledgements

We thank the members of Immunogenomics laboratory at National Institute of Immunology (NII) for useful discussions and suggestions.

## Funding

This work was supported by the DBT/Wellcome Trust India Alliance Fellowship (grant number IA/I/18/2/504012) awarded to VSP and DBT-NII core funding to VSP. RK is supported by DBT-JRF (Department of Biotechnology-Junior Research Fellowship) for his PhD.

## Contributions

RK and VSP conceived, designed, and executed the study. RK, KS and SS performed and analysed the experiments under the supervision of VSP, Genomics data was analysed by RK and SC under the supervision of GAA and VSP, AS supervised patient recruitment and sample collection, RK and VSP wrote the manuscript, which was further edited and approved by all authors.

## Ethics declaration

### Ethics approval and consent to participate

Approval for the use of human material for research was obtained from the institutional human ethics committees from both National Institute of Immunology, New Delhi and Vardhman Mahavir Medical College (VMMC) and Safdarjung Hospital, New Delhi, India. The donors consented for use of the plasma and buffy coat samples for research.

## Competing interests

The authors declare that they have no competing interests.

## Additional files

## Additional file 2: Supplementary Tables

**Table S1.** Details of cells from single-cell genomics experiments.

**Table S2:** Details of reagents used in the study

**Table S3:** Gene list used for enrichment analysis in the study

**Table S4.** Single-cell cluster wise enrichment of transcripts using MAST (log2FC>0.25; adj pval <0.05)

**Table S5:** TCR clonotype details of hCMV-reactive single-cells

**Fig. S1.**
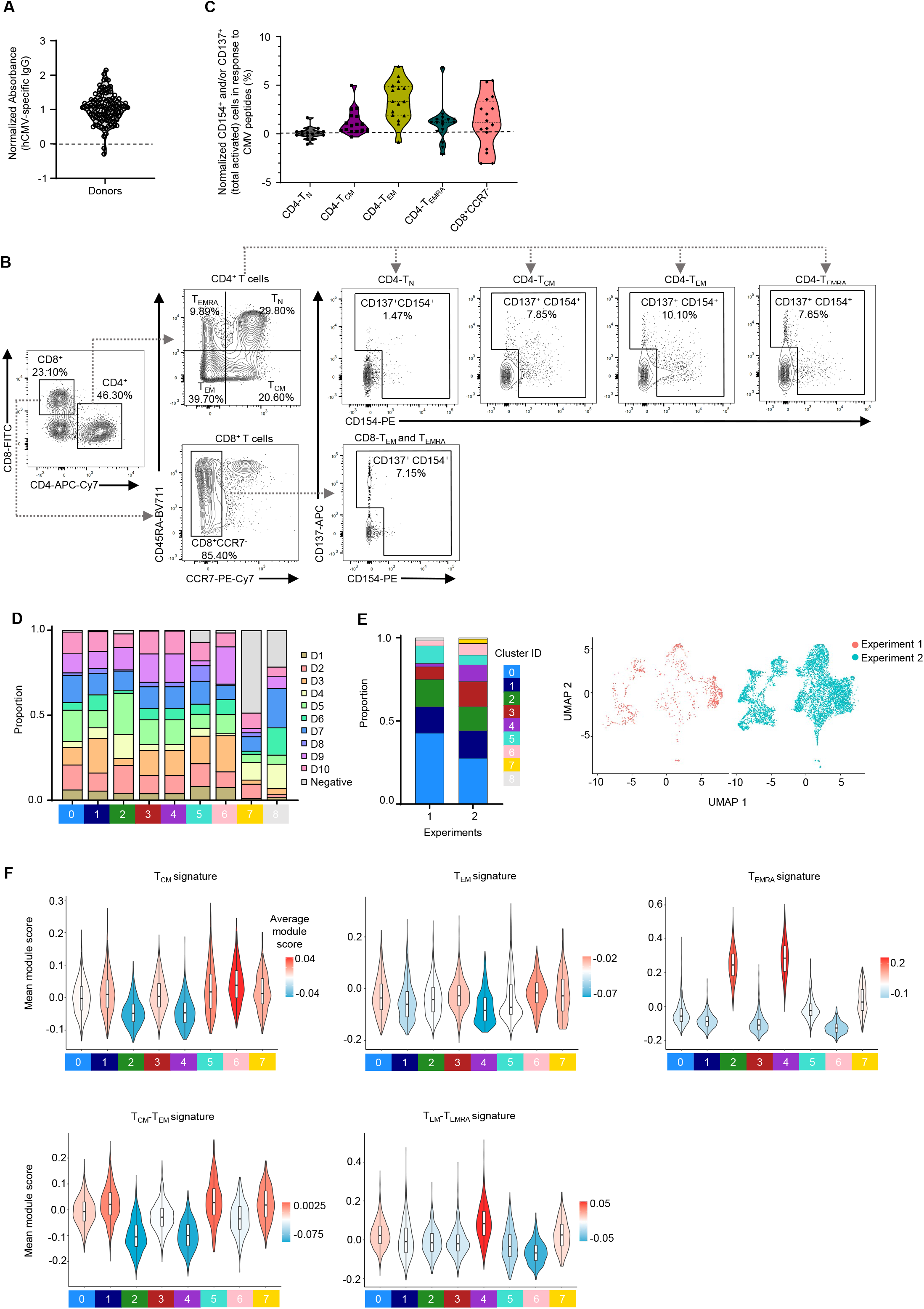
Single-cell RNA-Seq of hCMV-specific cells. (A) Graph shows the hCMV-specific IgG positivity (normalized absorbance) in 126 human donors. (B) Contour plots showing the gating strategy used to analyse the activated T cells in the indicated subsets upon stimulation of the PBMCs with hCMV peptide pools. (C) Violin plot shows the normalized T cell reactivity to hCMV-specific peptide pools compared to unstimulated control. Donors above the horizontal dotted line for memory compartments were chosen for further single-cell genomics studies (10 out of 16). (C) Stacked bar graph shows the distribution of cells from different donors across 9 clusters. (D) Stacked bar graph (left) and the 2D UMAPs (right) show the distribution of cells across 9 clusters from 2 different experiments/batch. Experiment 1 included 3 donors (D1, D2 and D7) and experiment 2 included all 10 donors. (E) Violin plots show the module enrichment score for T_CM_, T_EM_, T_EMRA,_ T_CM_-T_EM_, and T_EM_ -T_EMRA_ specific gene sets across 8 clusters. Colour scale indicates average module score. Clusters with <1% of the total cells are not shown (cluster 8=56 cells).

**Fig. S2.**
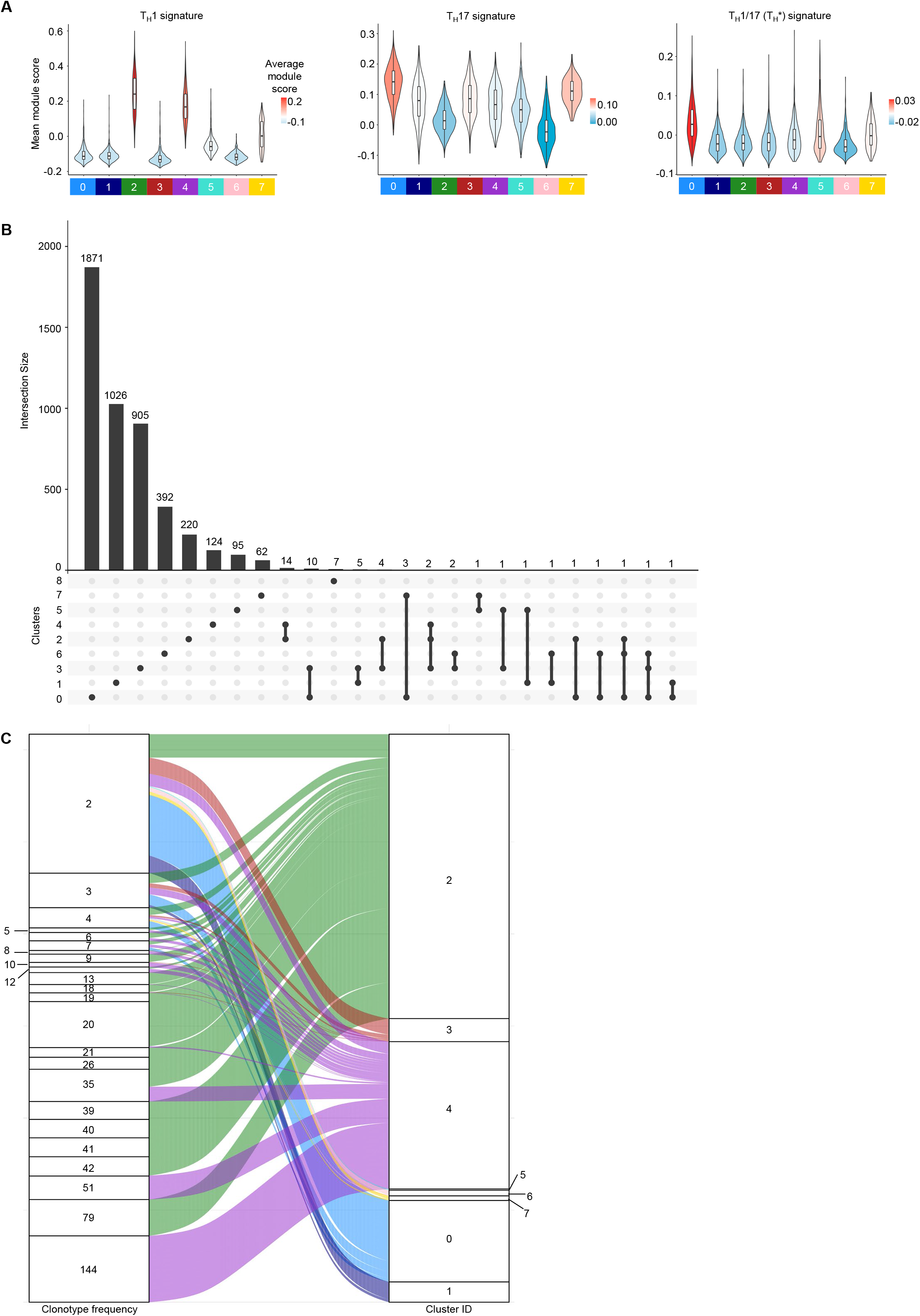
Functional characterization and TCR analysis of hCMV-specific T cells. (A) Violin plots show the module enrichment score for T_H_1, T_H_17, and T_H_1/17 subset-specific gene sets across 8 clusters. Colour scale indicates average module score. Clusters with <1% of the total cells are not shown (cluster 8=56 cells). (B) UpSet plot shows the TCR clonotypes (frequency on y-axis) and sharing of the clonotypes (bottom, connected dots) across the 9 clusters. (C) Alluvial plot shows the clonally expanded cells (1234 cells) and their distribution and sharing across different clusters. Each connecting line represents a single cell.

**Fig. S3.**
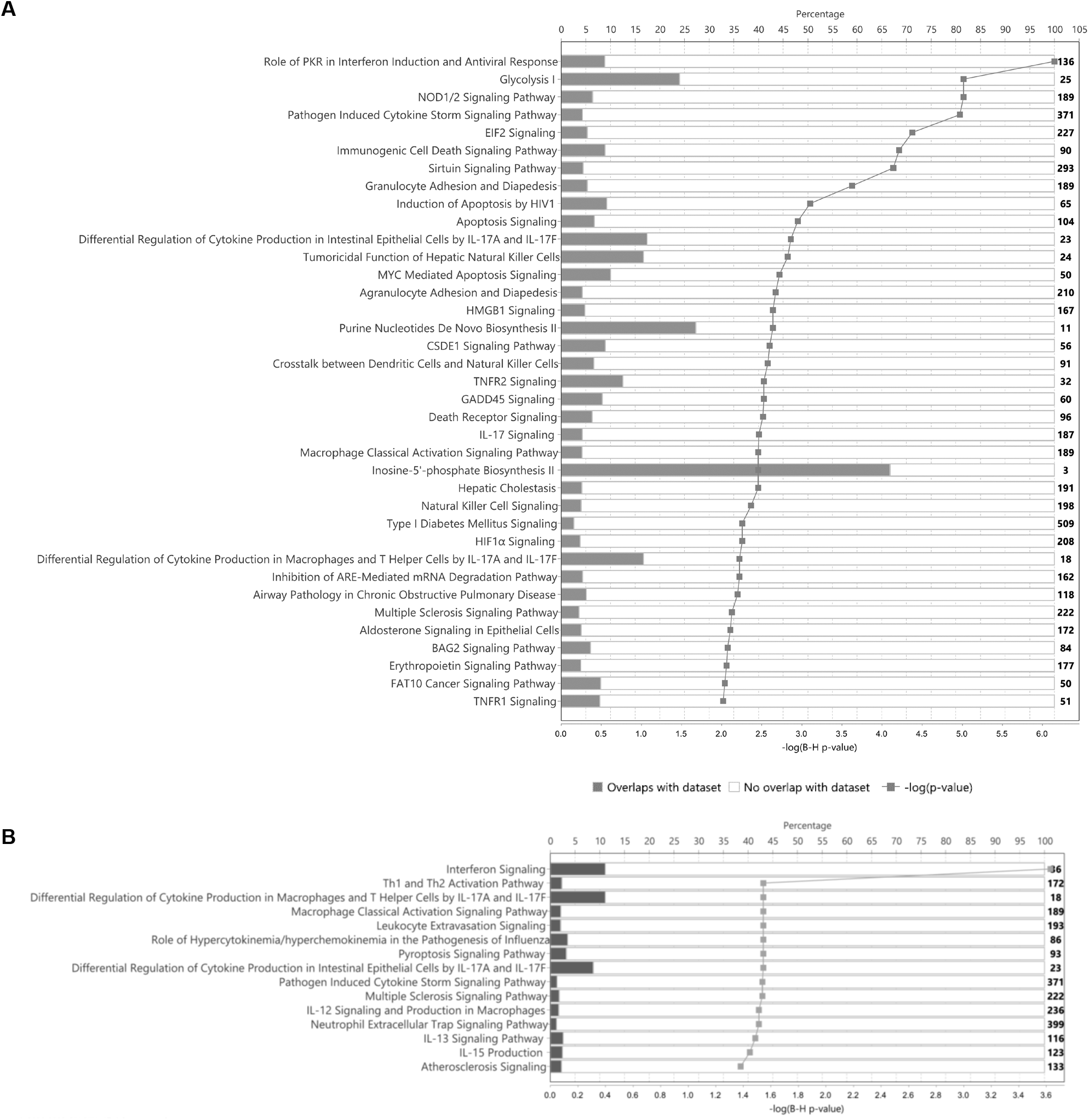
Pathways enriched in hCMV-specific cytotoxic clusters. (A) Pathway enrichment analysis using Ingenuity Pathway Analysis (IPA) (Qiagen) shows enrichment of pathways in cluster 2 compared to cluster 4. (B) Pathway enrichment analysis using IPA shows enrichment of pathways in cluster 4 compared to cluster 2.

## Notes

### Competing Interest Statement

The authors have declared no competing interest.

### Summary of Updates

Have uploaded a wrong pdf. Hence uploading the correct one here.

